# EGGS: Empirical Genotype Generalizer for Samples

**DOI:** 10.1101/2025.10.16.682896

**Authors:** T. Quinn Smith, Amatur Rahman, Zachary A. Szpiech

## Abstract

**Summary:** We introduce Empirical Genotype Generalizer for Samples (EGGS) which accepts empirical genotypes with missing data and replicates the distribution of missing genotypes along the empirical segment in other replicates. The empirical segment must have a number of sites less than the replicate. In addition, EGGS can remove phase, remove polarization, simulate deamination, simulate sequencing error, create pseudohaploids, and convert between Variant Call Format (VCF), *ms*-style replicates, and EIGENSTRAT/ANCESTRYMAP. When producing VCF files, EGGS is not limited to biallelic sites and assumes all samples are diploid.

**Availability and Implementation:** EGGS is written in the C programming language. Precompiled executables, source code, the manual, and the analysis conducted in the paper are available at https://github.com/TQ-Smith/EGGS

## 1 Introduction

Recent advantages in both forward and backward in-time simulations have made it possible to generate synthetic genotypes under varying evolutionary assumptions and complicated demographic scenarios [1, 2, 3, 4]. Such simulations are essential for rigorously testing computational methods, and more recently, training machine learning models [5, 6, 7, 8]. In addition, simulations are often used to test hypotheses concerning the evolutionary histories of human and nonhuman species [6, 9]. Simulated samples are reported under idyllic conditions where each genotype is phased, and in the case of biallelic sites, each site has a known ancestral allele. Simulated genotypes are devoid of technical artifacts that introduce variant uncertainty, such as missing alleles and deamination. Empirical data are often error-prone and violate the assumptions of simulated data. Ignoring this discrepancy in data quality can potentially lead to erroneous inference and unreliable results. Robust inference in the presence of low-quality genotypes is of particular interest when studying ancient DNA (aDNA) [10, 11]. This has motivated the development of methods to account for genotype uncertainty [12, 13, 14, 15, 16, 17].

Missing alleles are caused by compounding sources, including sample quality, sequencing technology, and data processing. The inherent stochasticity and complexity of such error are randomly introduced into simulated data [18]. This approach assumes that the proportion of missing samples at each site viewed in aggregate reflects the underlying structure of missing genotypes. However, a cumulative summary of missing sites for each sample ignores stretches of missing genotypes across the empirical segment, especially in regions of low complexity. In addition, the random modeling of missing data using a predefined distribution may inaccurately capture the empirical distribution. These difficulties are often exacerbated in non-model organisms lacking a high-quality reference genome [19]. Several recent studies introduce missingness blocks that correspond to regions of low mappability and variant callability [20, 21]. Other current methods that introduce missing genotypes require a FASTA file [22], ignore the position of the missing sites within the simulated segment, are restricted to specific simulation frameworks, simulate error prone reads directly [23], or are not generalizable to any supplied Variant Call Format (VCF) file [24].

We introduce Empirical Genotype Generalizer for Samples (EGGS) which extracts the underlying distribution and tracks of missing sites across a genomic region and introduces similar patterns of missing genotypes in replicates. To avoid confusion, we define a missing genotype at a locus where a sample is missing both alleles. In a VCF, this is denoted with “./.”. EGGS works on replicates with a number of sites less than the empirical segment and any number of samples. We demonstrate the use of EGGS and illustrate its ability to capture realistic patterns of missing genotypes for a simulated set of samples.

## 2 Materials and Methods

### 2.1 Replicating Missing Sites

Consider *S* samples genotyped at *N* sites. Our goal is to introduce the structure of missing genotypes in a synthetic replicate. Let *T* be the number of samples in the synthetic replicate, where the *T* samples are genotyped at *M < N* sites. *T* is not restricted by *S*. A missing genotype is a site where the sample is missing both alleles. We partition the *N* sites into *M* blocks. Each block contains 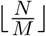 sites, except the first *N* (mod*M*) blocks which contain 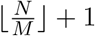 sites.

For each sample 1 ≤ *i* ≤ *S*, we calculate the average number of missing genotypes within a block. This quantity is denoted by *p*_*ij*_, where 1 ≤ *j* ≤ *M* . We introduce missing genotypes into the synthetic replicate by repeating the following process for each of the *T* samples: For each site 1 ≤ *j* ≤ *M*, randomly choose one of the *S* samples *s* ∼ *Uniform*(1, …, *S*). We label the site as missing with probability *p*_*sj*_.

Representing missing sites in *M < N* sites forces some amount of information to be lost. By partitioning the *N* sites into *M* blocks and summarizing the average number of missing genotypes for each sample within each block, we are capturing the larger trends of missing genotypes, and essentially, compressing the pattern of missing genotypes. The resampling procedure allows the missing genotypes to be recreated among many synthetic samples. Our method allows empirical segments containing regions with large amounts of missingness to be recreated in smaller sized synthetic replicates. This is analogous to how diploS/HIC introduces missing genotypes in VCF files [22]. Instead of using two VCF files, diploS/HIC partitions a FASTA given a VCF file and labels the site as missing for all samples depending on the current FASTA partition’s ‘N’ content [22].

### 2.2 Implementation

EGGS accepts variants in VCF [25] or *ms*-style formatting [26]. In addition, we provide an option for the conversion from EIGENSTRAT/ANCESTRYMAP format, the format used by The Allen Ancient DNA Resource (AADR), to VCF [27, 28]. In the case of multiple *ms*-style replicates as input, replicates are split to separate VCF files. We provide an option that splits each lineage to a separate sample set. This allows each lineage in an *ms*-style replicate or in a VCF file to be treated as its own sample in the resulting VCF. When creating VCF files, EGGS produces diploid samples.

To remove phase, EGGS provides two options. The first option switches the left and right allele at each genotype with equal probability. The second option places the smaller-index allele first in the genotype. In the case of VCF output, each unphased genotype is marked with ‘/’ as opposed to ‘|’. Computational methods that rely on phase typically ignore the VCF specification [25] and rely on the position of the left and right allele at a genotype. Therefore, changing ‘|’ to ‘/’ is not sufficient to erase phase information. The reference allele at each site is typically treated as the ancestral allele, which in turn, is used as the outgroup’s state in many analyses. The choice of the ancestral allele is known to affect inference [29]. Treating an allele as the ancestral is known as polarization. To remove polarization from biallelic sites, EGGS swaps the ancestral and derived alleles for all samples’ alleles by a user supplied probability. EGGS can introduce missing genotypes randomly according to the approach of Pandey et al. [18]. The proportion of samples with both alleles missing is calculated per site. The mean and standard deviation of the proportion of missing genotypes is calculated over all sites and is used to parameterize a *β*-distribution. The *β*-distribution is used to randomly choose the proportion of missing samples at each site in the supplied genotypes. The user can choose to directly give the mean and standard deviation to parameterize the *β*-distribution or a VCF from which the values are calculated.

Another complication commonly arising in working with aDNA is deamination. Deamination is a type of damage, when usually, a cytosine is incorrectly read as a thymine during sequencing [11]. Deamination is simulated according to Harney et al. [30]. The user supplies two values: the probability that a site is a transition and the probability that a called reference allele will deaminate to the alternative allele. Deamination is synthetically introduced into the dataset by applying the above to every sample’s alleles at all site. An additional complication arising with aDNA is the presence of pseudohaploids. The low quality of aDNA makes confidently calling heterozygotes difficult. Instead, at sites with mapped reads showing multiple alleles, one allele is randomly chosen [18]. Pseudohaploids are created by randomly choosing each sample to be homozygous for either the reference or alternative allele with equal probability at sites where the sample contains both the reference and alternative allele. For sites where one of the alleles are missing, the non-missing allele is made homozygous.

Modern-day genomes are less degraded compared to aDNA and contain much lower error rates on next-generation sequencing machines [31]. We introduce sequencing error by switching the state of every sample’s called allele at a biallelic site with a user supplied probability. Finally, summary statistics regarding missing genotypes can be calculated similar to those implemented in PLINK [32]. EGGS can produce *ms*-style output when missing data is not introduced. Some options cannot be used together to avoid ambiguity in precedence. For example, we do not allow sequencing error to be introduced when deamination is being introduced. This is described in the manual. EGGS is written in the C programming language to enable fast processing of thousands of replicates. Precompiled executables, source code, and manual are available at https://github.com/TQ-Smith/EGGS.

## 3 Results

We used EGGS to convert the Allen Ancient DNA Resource (AADR) [27] to VCF, and we extracted chromosome 1 of the 217 samples exclusively from Mathieson et al. [33] with bcftools [34]. This resulted in 93166 sites. The sites had a mean missingness of 0.55 and a standard deviation of 0.21. We refer to this set as the ancient samples.

Using *msprime* [2], we simulated 200 diploid samples on segments ranging from 1Mb to 10Mb in 1Mb increments. The segments were generated with the effective population size *Ne* = 10, 000, mutation rate *µ* = 1.25 × 10^*−*8^ per base pair, and the recombination rate *ρ* = 1 × 10^*−*8^ per base pair. We generated a range of segment lengths because the number of segregating sites in each replicate is the important variable when evaluating EGGS. We ran EGGS 10 times for each replicate introducing missing sites with EGGS’s method and using the *β*-distribution method. At each site, we average the proportion of samples missing for each method across the 10 runs. The average proportion of samples missing at each site for each segment length is used for comparison with the ancient samples.

Dynamic time warping (DTW) is a signal processing technique used to compare two signals of different lengths assuming that one signal is a slowed down or sped up version of the other. DTW is often used to evaluate compression or interpolation methods and is analogous to sequence alignment in bioinformatics [35]. We use DTW to evaluate our method. We compute the proportion of missing samples at each of the 93166 sites in our ancient samples. We treat this as a signal and use DTW to compare it to the average signal produced by our simulated replicates. DTW computes the Euclidean distance between two aligned time points. A lower value indicates that the two signals are more similar. We purposefully omit the units of this value as the interpretation is difficult in this context. The results of the DTW for the replicates is shown in Table 1.

**Table 1:** Results of DTW. The first column is the length of the simulated segment in megabases. The second column is the number of segregating sites in the simulated segment. The third column is the minimal number of segregating sites in the block. The fourth column is the distance as calculated by the DTW between the missingness in the empirical segment and the missingness introduced in the simulated replicate using the *β*-distribution. The fifth column is the distance as calculated by the DTW between the missingness in the empirical segment and the missingness introduced in the simulated replicate using EGGS’s method. For the *β*-distribution and EGGS’s method, the DTW was calculated with the average proportion of missing samples at each site in the simulated segment over 10 runs.

| Length of Segment (Mb) | Number of Segregating Sites | Block Size | β-Distribution | EGGS |
| --- | --- | --- | --- | --- |
| 1 | 3327 | 28 | 57.49 | 57.38 |
| 2 | 6612 | 14 | 55.79 | 55.02 |
| 3 | 9657 | 9 | 54.57 | 53.09 |
| 4 | 12659 | 7 | 53.46 | 51.39 |
| 5 | 16468 | 5 | 52.51 | 49.2 |
| 6 | 19930 | 4 | 51.53 | 47.33 |
| 7 | 23012 | 4 | 51.08 | 45.67 |
| 8 | 25682 | 3 | 50.45 | 44.29 |
| 9 | 29391 | 3 | 49.84 | 42.39 |
| 10 | 33023 | 2 | 49.27 | 40.92 |

For all segment lengths, we see that EGGS’s method captures the proportion of missing samples along the empirical chromosome better than the approach of the *β*-distribution. The discrepancy between the two methods is smaller for shorter segments and becomes larger for longer segments. This suggests that introducing missing data into shorter segments according to a *β*-distribution may be sufficient when missing data is not a concern for downstream analysis; however, as the number of segregating sites in the simulated segment approaches the number of segregating sites in the empirical segment, EGGS’s method better realizes fluctuations in the proportion of missing samples.

## 4 Discussion

We introduced Empirical Genotype Generalizer for Samples (EGGS) that artificially removes genotypes in patterns similar to empirical data as opposed to treating the distribution of missing data across a segment as purely random. Combining EGGS’s ability to remove genotypes and other evolutionary assumptions, we can quickly make simulations more realistic, enabling more robust downstream analyses.

EGGS’s method to replicate missing genotype patterns becomes limited when there is a large difference in the number of loci between the set of variation containing the patterns to replicate and the set of variation in which missing genotypes will be introduced. Such a discrepancy between the number of loci causes the resolution of EGGS’s stratification method to decline. In addition, we expect that in situations where the rate of missing genotypes is high and the segment is small (less than 1Mb), EGGS’s approach will not yield significantly different results than modeling missingness with a beta-distribution. We also assume that adjacent missing sites for samples are not correlated. While this assumption is justified for aDNA [18], this is not justified for less error prone data. In such a case, our approach could be replaced with more sophisticated signal processing techniques and offers a path for future work [36]. Finally, we note that our approach could be adapted to replicating other characteristics of genotype uncertainty, such as homozygosity and ascertainment [37, 38, 39].

## 5 Acknowledgments

The authors would like to thank Abigail N. Sequeira, Ana V Leon-Apodaca, Anna Maria Calderon, and three reviewers for helpful suggestions. Computations for this research were performed using the Pennsylvania State University’s Institute for Computational Data Sciences’ Roar supercomputer. This work was supported by the National Institute of General Medical Sciences of the National Institutes of Health award number R35GM146926 (ZAS).

